# Successful coral microbiome transplant from high to low heat tolerant corals requires antibiotic pretreatment

**DOI:** 10.64898/2026.08.28.747763

**Authors:** Lindsey K. Deignan, Clarence W. H. Sim, Keay Hoon Pwa, Rebecca J. Case

## Abstract

Microbiome transplantation, used to treat human disease, can enhance thermal and pathogen resilience in bleaching-susceptible corals via coral microbiome transplantation (CMT), though success is donor- and recipient-dependent. In this study, less thermally tolerant *Pachyseris speciosa* fragments were exposed to ciprofloxacin or an antibiotic cocktail for 24 h, then received CMT from the thermally tolerant *Acropora millepora* from Singapore’s turbid reef system. Alpha diversity increased only in antibiotic-treated, CMT fragments, demonstrating that antibiotic-induced dysbiosis enhanced bacterial uptake. Coral microbiome assemblage shifted significantly at 1 and 10 d, regardless of antibiotic treatment or *Acropora* inoculum. Antibiotic-induced dysbiosis did not enhance uptake of donor’s core ASVs (e.g., *Endozoicomonas* spp.). However, early uptake favoured potential pathogens like *Vibrio* spp., while longer inoculation allowed for uptake of unculturable environmental taxa.

Our approach of using antibiotic pretreatment followed by whole microbiome transplant parallels human faecal microbiota transplantation to restore gut health.

## Introduction

A coral holobiont consists of the coral host, endosymbiotic dinoflagellates, and the associated microorganisms of the microbiome, primarily bacteria, archaea, fungi and viruses^1–3^. The microbiome supports the coral host through a number of functions, including pathogen deterrence and facilitating metabolic pathways, such as nitrogen fixation and sulphur cycling^4^. The microbiome can help mediate the response of the coral holobiont to environmental perturbations, thus enhancing the resilience of the coral^4–7^. While the stress response of coral microbiomes is species-specific^6^, those species with more flexible microbiomes, with a lower proportion of co-evolved species, are thought to possess more resilience potential^5,8,9^.

The ‘beneficial microorganisms for corals’ (BMC) hypothesis proposes that the coral microbiome can be manipulated and restructured to increase coral adaptability to environmental stressors^3,10^, particularly those associated with global climate change.

Experimental trials have shown that the uptake of potential BMC is effective in conferring useful traits, namely heat or pollution tolerance and disease resistance^11–15^. In addition to inoculation of BMC, the direct inoculation of whole microbiome from tissue homogenate, or coral microbiome transplantation (CMT), has effectively conferred environmental stress resilience, when heat-tolerant coral microbiomes were inoculated onto heat-susceptible individuals^16^. The ability to confer heat-tolerance of BMCs that resist culturing, and avoid the time intensive screening and cultivation of individual BMC makes CMT an appealing method for rapid deployment.

As coral microbiome manipulation proves itself as an effective tool to temporarily enhance coral stress tolerance, there is growing acknowledgement of the need to develop approaches that can elicit a more sustained effect, to avoid the requirement of repeat inoculations^17^. One potential strategy to prolong retention of a manipulated microbiome is to disturb and suppress the existing microbial community with antibiotic treatment before inoculation, thereby providing spatial (habitat) and nutrient resources for new microbes to colonize and lowering competition with resident microbes. Antibiotic treatment of corals can temporarily disrupt the microbiome, termed dysbiosis, without impacting key drivers of coral health, like photosynthetic yield or host tissue protein content^18–20^, if corals are not exposed to major environmental stress^19,21^. Dysbiosis provides a window of time to administer CMT immediately following antibiotic treatment after the native prokaryotic community has been suppressed.

Our study took place in Singapore, an island nation located west of the coral triangle, where coral reefs occur within a highly urbanized seaport. Despite widespread coral bleaching events reported in recent years^22,23^, Singaporean coral reefs remain resilient, surviving fast-developing seascapes, high turbidity and sedimentation rates, and increasing temperatures^24^. The plating coral *Pachyseris speciosa* is found abundantly on Singapore’s reefs, typically occurring in the low light environments of its lower reef^24–27^. It is a high risk species for bleaching due to its sensitivity to elevated temperatures^28^. *P. speciosa* exists in two distinct lineages within Singapore^27^; therefore, we refer to it generically as *Pachyseris* hereafter. Conversely, *Acropora millepora* (referred as *Acropora* hereafter) has been observed locally to be resistant to bleaching during periods of high temperature where *Pachyseris* faired poorly^25^.

We conducted a mesocosm-based CMT experiment to test if pre-treating *Pachyseris* with a single or mixed cocktail of antibiotics, to suppress its native bacterial community, increases the uptake of ASVs from the heat-tolerant *Acropora’s* microbiome. Specifically, we aimed to:

1. determine whether antibiotic-mediated dysbiosis of native bacterial microbiome enhances uptake of ASVs from the donor *Acropora* microbiome to the recipient, *Pachyseris*.
2. assess whether there is prolonged retention of core ASVs from donor to recipient corals.

## Results

To test whether antibiotic induced dysbiosis and suppression of the native microbiome improves the uptake and persistence of a transplanted coral microbiome, we characterized the bacterial community using 16S rDNA metabarcoding of *Pachyseris* fragments across antibiotic treatments (control with no antibiotic, antibiotic cocktail and ciprofloxacin), with and without subsequent inoculation with *Acropora’s* whole microbiome (Figure 1).

**Figure 1.**
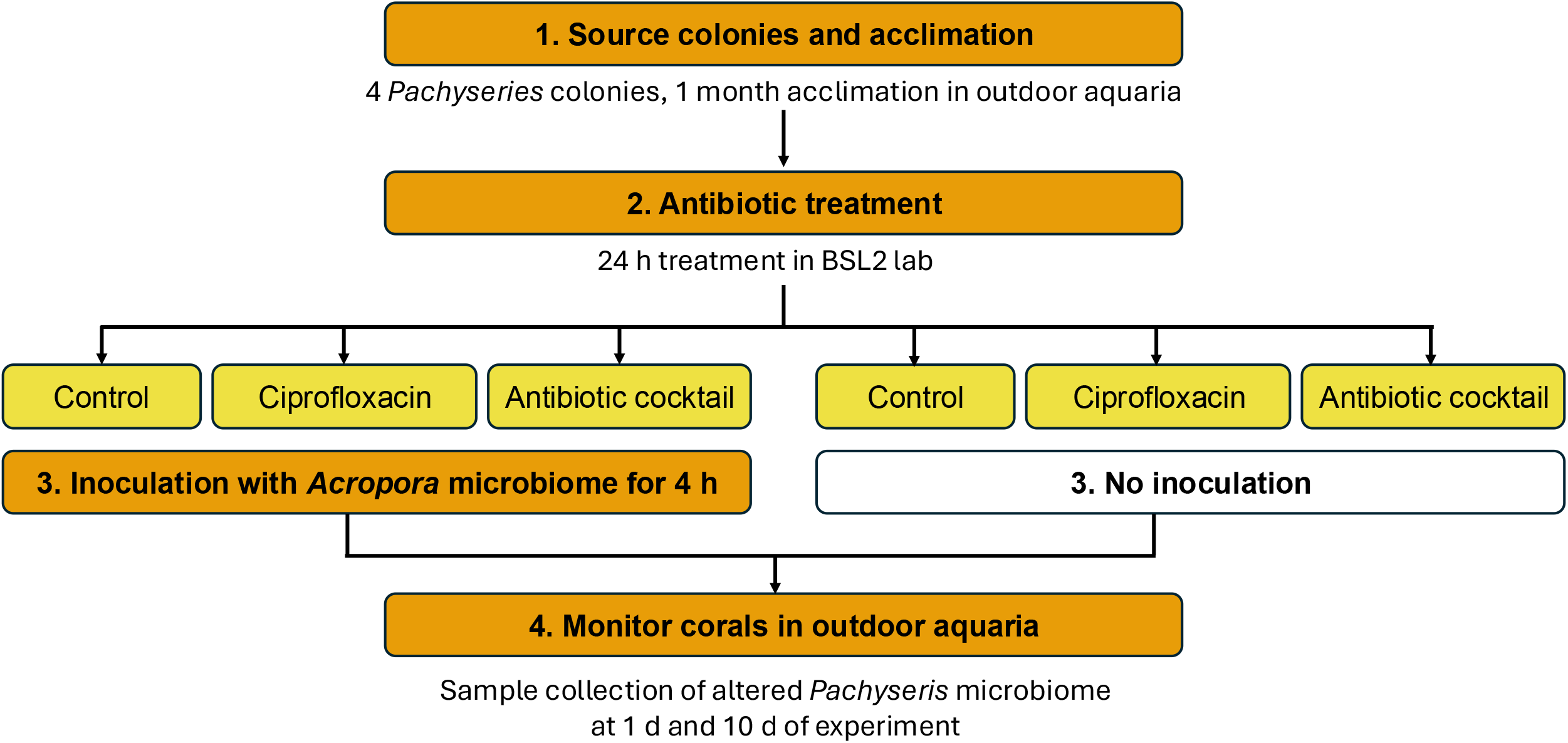
Experimental design part 1: Initial collection and preparation began with collection of four *Pachyseris* colonies, followed by a one month acclimation period in outdoor aquaria. Part 2: coral fragments subjected to 24 h antibiotic cocktail treatment (BSL2), split across three experimental conditions: an untreated control group, a ciprofloxacin treatment group, and an antibiotic cocktail group. Part 3: Following antibiotic exposure, half of the fragments from each treatment underwent a 4 h *Acropora* inoculation. Part 4: All groups were returned to individual outdoor aquaria for monitoring, with triplicate sampling conducted on 1 and 10 d post-treatment.

### *Pachyseris*’ microbiome requires antibiotic treatment for *Acropora* CMT to shift its diversity and assemblage

*Pachyseris* fragments not inoculated with *Acropora* microbiome did not experience any significant temporal impacts on alpha diversity, represented by Shannon index, regardless of antibiotic treatment (Figure 2A, Table S1). However, we note that ciprofloxacin led to a smaller variability in diversity compared to the antibiotic cocktail and control (Figure 2A, Table S1). At 10 d, both antibiotic treatments were significantly different (t-test, p < 0.05) to the control and each other, but no treatment (control, antibiotic cocktail, ciprofloxacin) differentiated from inoculated samples within the same treatment at either 1 or 10 d. In contrast, samples inoculated with *Acropora* microbiome experienced increased alpha diversity at 10 d, with statistically significant (t-test, p < 0.05) increases for those treated with either antibiotic treatment (Figure 2B, Table S1).

**Figure 2:**
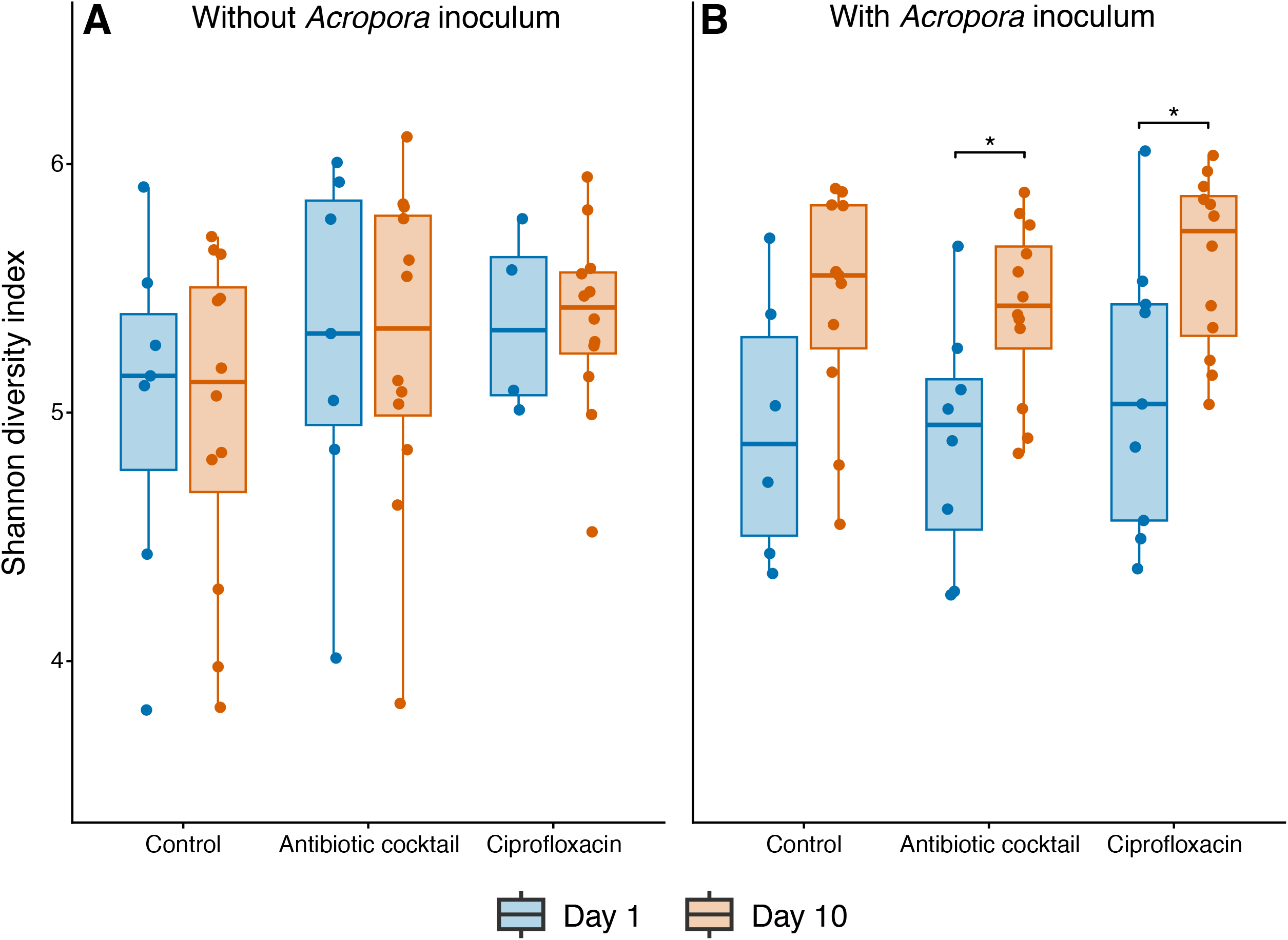
Shannon diversity index of the *Pachyseris* microbiome at 1 and 10 d post- inoculation, across the three antibiotic treatments (Control, antibiotic cocktail, ciprofloxacin), for coral fragments (A) without and (B) with *Acropora* microbiome inoculation. Data points represent individual fragments. Asterisks (*) indicate a significant increase in Shannon diversity from 1 to 10 d of the experiment within a treatment (Welch’s two-sample t-test, Benjamini–Hochberg adjusted p < 0.05).

To resolve whether bacterial community turnover between 1 and 10 d occurred independently within each treatment and inoculation combination, separate Bray-Curtis NMDS ordinations and PERMANOVA tests were performed comparing 1 and 10 d within each treatment group, split by inoculation status (Figure 3). Community beta diversity differed significantly between 1 and 10 d in every treatment and inoculation combination (PERMANOVA, all p ≤ 0.003; Figure 3), a trend not observed in alpha diversity. For samples inoculated with *Acropora* microbiome, the communities converged to being more similar at 10 d compared to the larger spread of beta diversity at 1 d (Figure 3D-F) despite having higher alpha diversity at 10 d.

**Figure 3.**
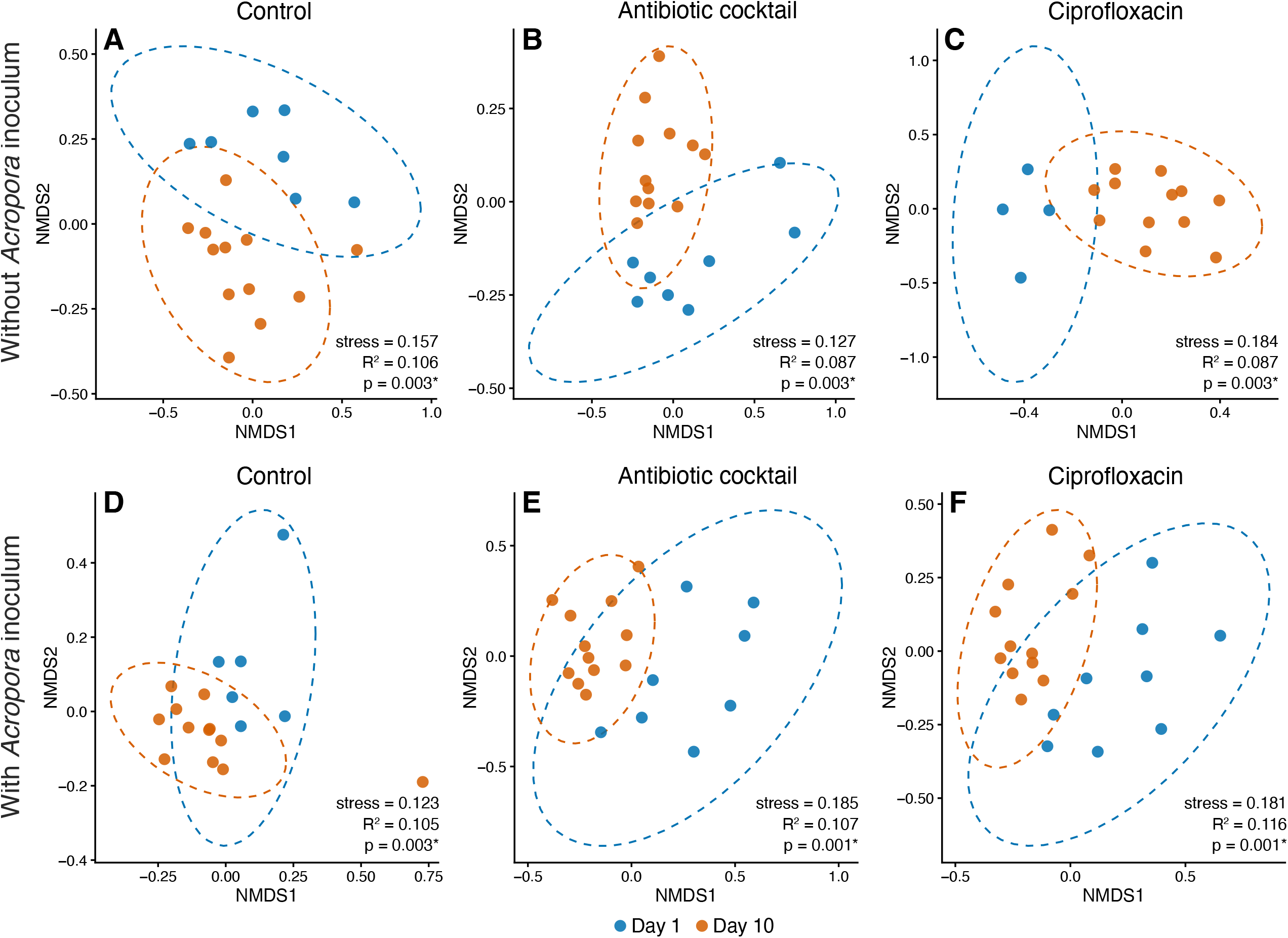
Non-metric multidimensional scaling (NMDS) ordinations of Bray-Curtis dissimilarities in the *Pachyseris* microbiome, comparing 1 and 10 d post-inoculation, shown separately for each antibiotic treatment (columns: control, antibiotic cocktail, ciprofloxacin) for fragments (A-C) without and (D-F) with *Acropora* microbiome inoculum. Dashed ellipses show 95% confidence regions (multivariate t-distribution) for each timepoint.

The effect of antibiotic treatment on community turnover at 1 d and 10 d were also resolved with Bray-Curtis NMDS and PERMANOVA tests (Figure 4, Table 1). Pairwise comparisons with the control showed significant shifts in community assemblage for both the antibiotic treatments of cocktail and ciprofloxacin, at both time points <u>Uptake of specific ASVs from *Acropora* CMT</u>

**Figure 4.**
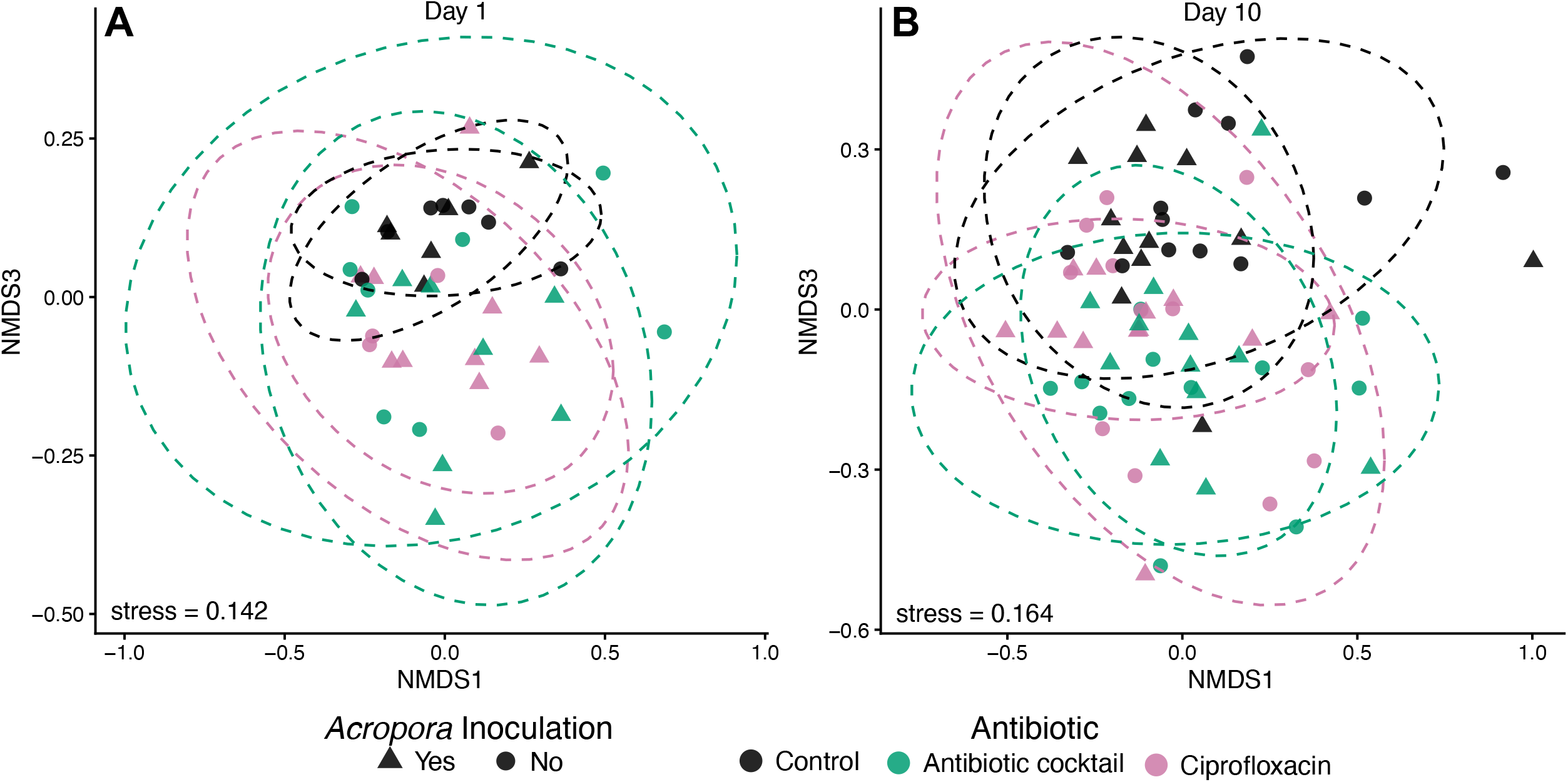
Non-metric multidimensional scaling (NMDS) ordinations of Bray–Curtis dissimilarities in the *Pachyseris* microbiome, in (A) 1 d and (B) 10 d post-inoculation, shown separately for each antibiotic treatment (colors), with (triangle) and without (circles) *Acropora* microbiome inoculum. Dashed ellipses show 95% confidence regions (multivariate t-distribution) for each treatment group. Pairwise PERMANOVA results are listed in Table 1. Samples were ordinated on a three dimensional scale as two resulted in stress > 0.2.

**Table 1:** Pairwise PERMANOVA comparison between antibiotic groups corresponding to NMDS plots in Figure 4. Adjusted p-values < 0.05 are indicated by asterisks (*) and bold.

| Inoculation | Comparison | Day 1 |  | Day 10 |  |
| --- | --- | --- | --- | --- | --- |
|  |  | R<sup>2</sup> | adjusted p-value | R<sup>2</sup> | adjusted p-value |
| With<br>*Acropora*<br>microbiome | Ciprofloxacin vs Control | 0.104 | **0.042 \*** | 0.062 | **0.030 \*** |
|  | Antibiotic cocktail vs Control | 0.120 | **0.024 \*** | 0.061 | **0.030 \*** |
|  | Ciprofloxacin vs Antibiotic cocktail | 0.064 | 0.388 | 0.051 | 0.090 |
| No<br>inoculation | Ciprofloxacin vs Control | 0.125 | 0.072 | 0.070 | **0.006 \*** |
|  | Antibiotic cocktail vs Control | 0.107 | **0.0180 \*** | 0.086 | **0.006 \*** |
|  | Ciprofloxacin vs Antibiotic cocktail | 0.087 | 0.560 | 0.052 | 0.061 |

To determine which ASVs are core components of the *Acropora* microbiome, an Indicator Species Analysis (ISA) was used to compare *Acropora* microbiome samples to *Pachyseris* microbiome samples that did not undergo inoculation (control). The relative abundances of these indicator ASVs were then compared between uninoculated and inoculated treatment groups for detection of positive uptake of these ASVs by their new host, *Pachyseris*. There were 30 ASVs identified as highly significant (p < 0.01) indicators of *Acropora’s* microbiome, of which two were identified as *Endozoicomonas* spp. with high relative abundances within the microbiome (Figure 5A; Table S2). However, these *Endozoicomonas* were not established in any of the inoculated samples at 10 d (Figure 5B). The taxonomic identities of the 16 ASVs that were increased in inoculated *Pachyseris* samples include *Pyruvatibacter,* Methylotrophs, Alteromonaceae, *Shimia,* Rhodobacteraceae, *Fuerstia* and Planctomycetes.

**Figure 5.**
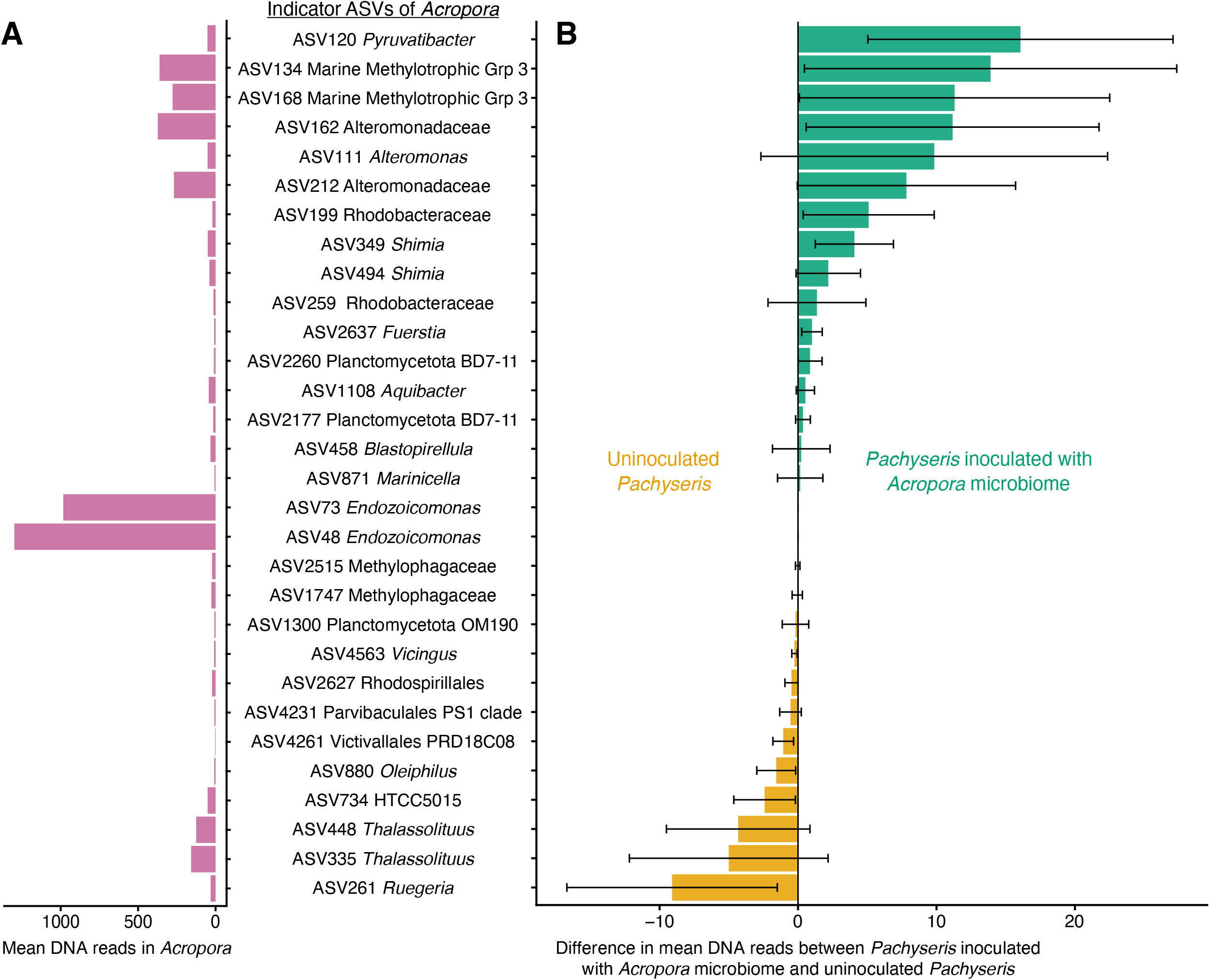
Differential abundance of 30 *Acropora*-associated indicator ASVs in *Pachyseris*, in descending order by relative enrichment in inoculated versus uninoculated fragments. (A) Mean number of DNA reads per ASV in donor *Acropora* tissue (n = 5). (B) Mean difference in DNA reads for the same ASVs between inoculated and uninoculated *Pachyseris* fragments with error bars showing ±1 SE. Blue bars: higher mean reads in inoculated fragments; orange bars: higher mean reads in uninoculated fragments. None of the differential abundances were statistically significant (all p > 0. 05).

ASVs shared between the *Acropora* microbiome inoculum and the inoculated treatment groups, but not found in uninoculated fragment groups, indicates the uptake of ASVs from CMT by the new host *Pachyseris*. To identify these ASVs, *UpSetR*^29^ analyses were performed. At both 1 and 10 d, the inoculum contained 79 ASVs that were not present in uninoculated *Pachyseris* samples (Figures 6 and 7). Of those 79 ASVs, at 1 d, two *Vibrio* ASVs were present in all three groups (control, antibiotic cocktail, ciprofloxacin) of antibiotic treated samples with the inoculum, as well as the donor 1 d *Acropora* microbiome (Figure 6; Table S3). An additional ASV from the Order Defluviicoccales (Proteobacteria) and *Staphylococcus* sp. ASV were also found in the antibiotic cocktail and ciprofloxacin treated samples inoculated with *Acropora* microbiome.

**Figure 6.**
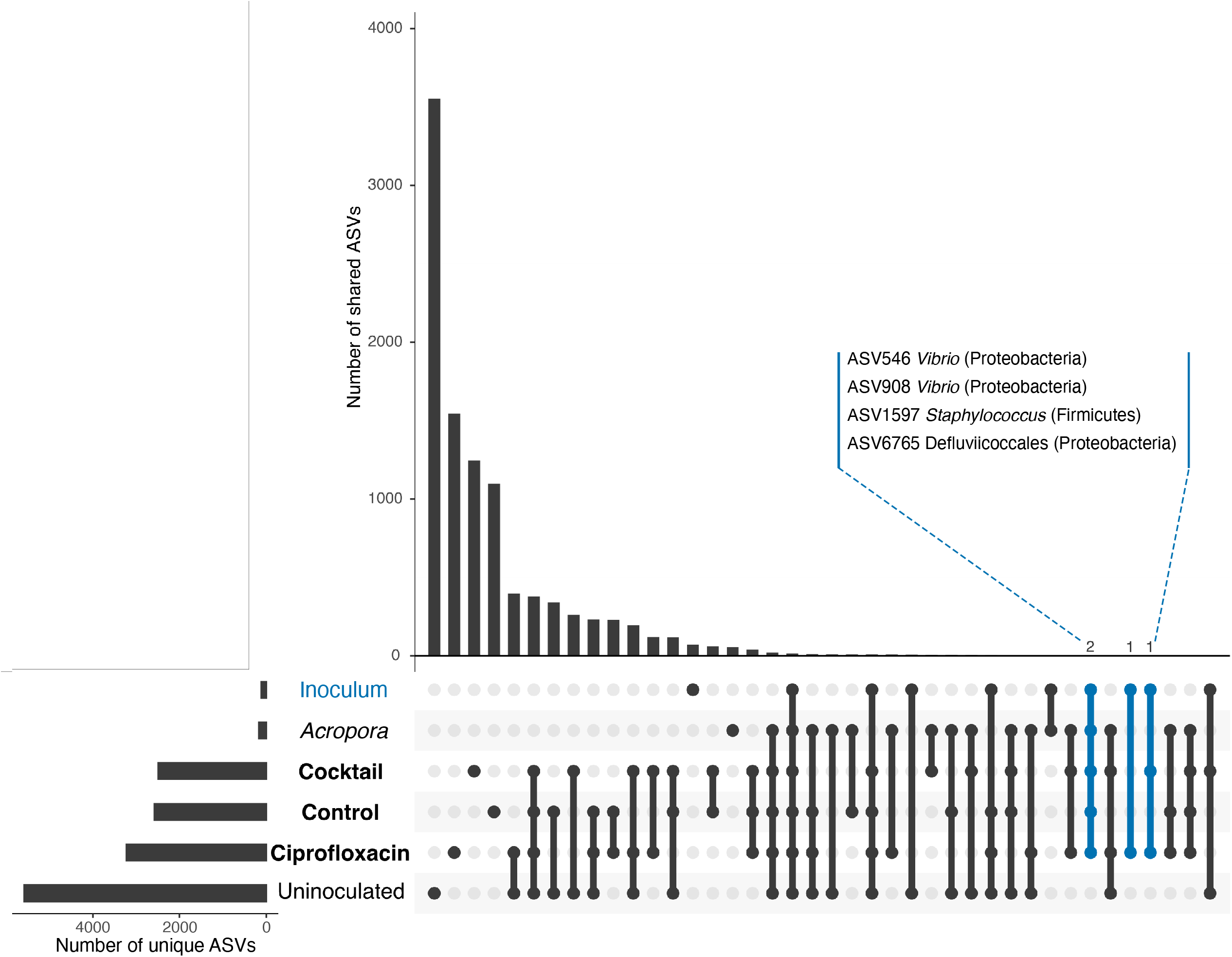
UpSet plot of shared and unique ASVs among sample groups at Day 1 post-inoculation: uninoculated *Pachyseris* fragments, *Pachyseris* fragments inoculated under each antibiotic treatment (labelled “Cocktail”, “Control”, “Ciprofloxacin”), donor *Acropora* coral tissue (“*Acropora*”), and the *Acropora*-derived inoculum suspension used for transplantation (“Inoculum” in blue). Horizontal bars (bottom left) show unique ASV richness per group; vertical bars show the number of ASVs shared among the groups indicated in the dot matrix below. Orange-highlighted intersections denote ASVs shared among inoculum-associated groups (must be absent from uninoculated fragments).

**Figure 7.**
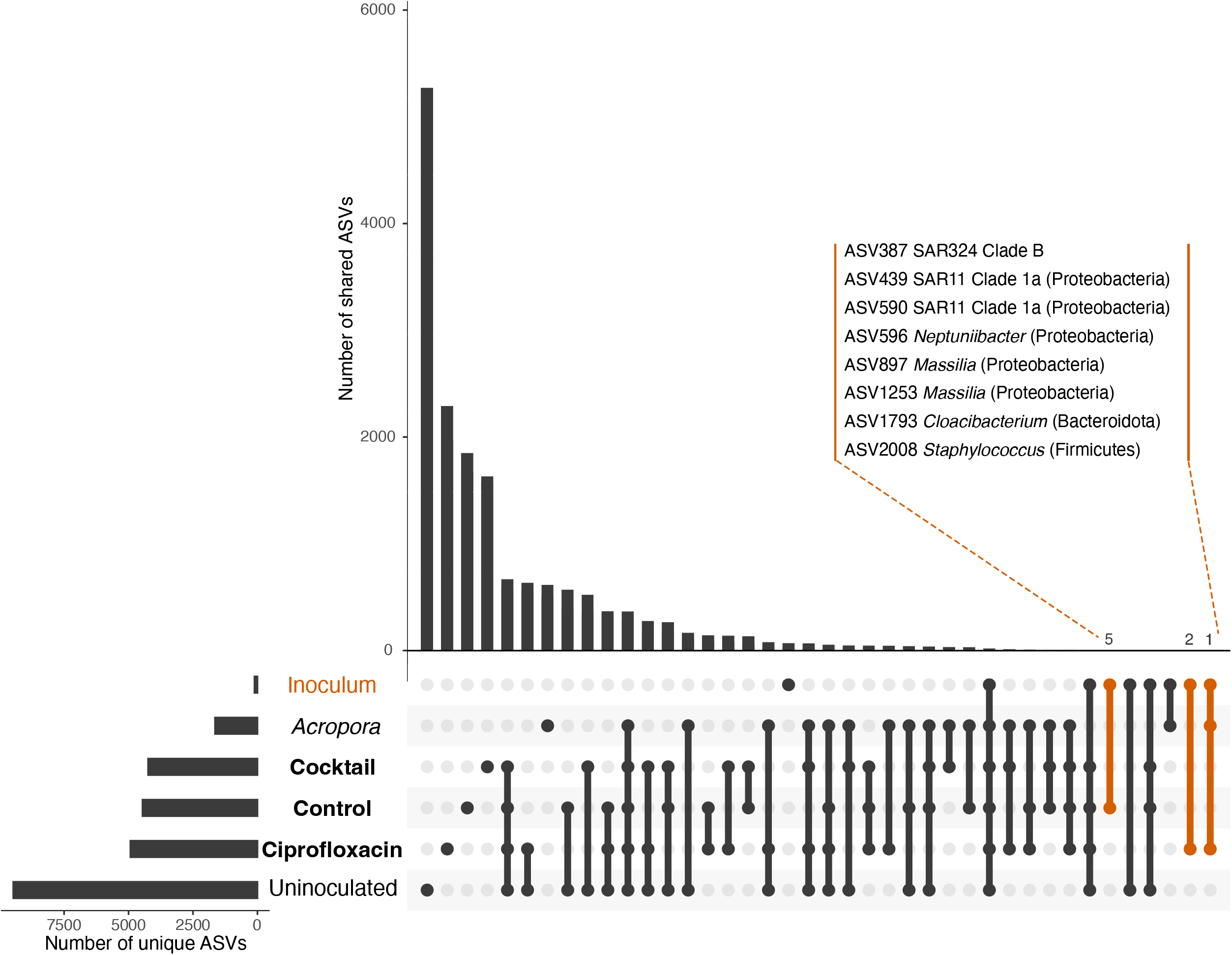
UpSet plot of shared and unique ASVs among sample groups at Day 10 post-inoculation: uninoculated *Pachyseris* fragments, *Pachyseris* fragments inoculated under each antibiotic treatment (labelled “Cocktail”, “Control”, “Ciprofloxacin”), donor *Acropora* coral tissue (“*Acropora*”), and the *Acropora*-derived inoculum suspension used for transplantation (“Inoculum” in orange). Horizontal bars (bottom left) show unique ASV richness per group; vertical bars show the number of ASVs shared among the groups indicated in the dot matrix below. Orange-highlighted intersections denote ASVs shared among inoculum-associated groups (must be absent from uninoculated fragments).

At 10 d, the control group without antibiotic treatment had five ASVs and while the ciprofloxacin treated group had three ASVs that were uniquely shared with the inoculum (Figure 7; Table S3). There were no ASVs uniquely present in the inoculum and the antibiotic cocktail treated group at 10 d. Strikingly, no same ASVs were detected as being recruited by *Pachyseris* at both 1 and 10 d (Figures 6 and 7; Table S3).

## Discussion

Intraspecific differences in stress tolerance occur naturally within a given location^30–32^, and Singapore’s equatorial turbid reefs provide natural stress gradients to support extended conspecific stress tolerance^31,33,34^ making assisted evolution (e.g. selective breeding, assisted gene flow, and epigenetic programming^35–38^) a potential avenue to produce thermally adapted corals. CMT is another potential approach for rapid impact on low stress tolerance corals. Coral microbiomes are species-specific^39^; thus, corals may be more susceptible to conspecific CMT. However, Doering, et al. ^16^ used CMT to impart temporary thermal stress tolerance in two corals: a *Pocillopora* sp. and *Porites* sp. suggestive that bacteria not co-evolved or specific to a coral host can move between coral species to extend a coral’s thermal range. In this study, using antibiotics to disrupt the host microbiome before inoculating the foreign microbiome was not enough to overcome the bacterial microbiome host specificity. However, CMT methodology to increase alpha diversity and shift beta diversity, without acquisition of donor ASVs with high relative abundance supports that idea that CMT of broad host range ASVs is possible from CMT.

### Antibiotic induced dysbiosis essential for CMT to increase bacterial diversity and restructure the bacterial community

Treatments that received both antibiotics and inoculum experienced a significant increase in alpha diversity over the course of the experiment (Figure 2B), indicative that a combinatorial treatment that employs antibiotic perturbation for CMT to increase the bacterial diversity within the holobiont, and that CMT alone did not increase bacterial diversity. The requirement of antibiotics for effective CMT supports that perturbation, or dysbiosis, is required for effective manipulation of the microbiome as perturbation opens niches and/or reduces interconnected relationships within the community, the foundation of stability for high diversity systems.

Ciprofloxacin-treated fragments showed less variability in diversity than those receiving the antibiotic treatment or CMT alone (no antibiotic treatment) (Figure 2A). As a single-compound fluoroquinolone, ciprofloxacin acts through the inhibition of bacterial DNA gyrase and topoisomerase IV, producing a singular selective pressure on the bacterial community. The antibiotic mixture combined four compounds (ampicillin, nalidixic acid, metronidazole and streptomycin) that act on distinct cellular targets, cell wall synthesis, DNA replication, anaerobic metabolism and protein synthesis, respectively^40^. Combinations of antibiotics with different modes of action can produce synergistic or antagonistic effects within a bacterium, and for the bacterial community, as four targets with additional combinatorial disturbance, make it difficult to predict antibiotic treatment outcomes, though presumably there is potential to cause broader perturbation across the bacterial community^41,42^.

### Is a single or cocktail antibiotic pretreatment, and time, required for effective CMT?

All treatments, and the controls, restructured their bacterial community within coral fragments from 1 to 10 d, so time is a consistent influence on the bacterial community structure (Figure 3), although not on alpha diversity (Figure 2). Both antibiotic treatments also significantly altered the bacterial community compared with untreated coral fragments at 1 and 10 d, (Figure 4) and so both time and antibiotics alter community structure as a predictable selective pressure. However, the antibiotic treatments were not significantly different to each other (Table 1, Figure 4), and so both a single and cocktail antibiotic strategy can be considered successful CMT strategies for perturbing the community as both result in a significant increase in diversity after CMT and a shift in bacterial assemblage.

Follow-up heat stress experiments are required to determine if thermal tolerance was conferred, however, the goal of this study is to determine if antibiotic pretreatment is required for CMT transplant to be effective, and if single or multiple antibiotics have a differential effectiveness in CMT impact on alpha and beta diversity.

### Uptake of specific donor ASVs via CMT

ASVs transferred from donor to recipient at 1 d included two *Vibrio* ASVs and one *Staphylococcus* ASV (Figure 6). Both genera include opportunistic pathogens of corals found in Singapore, often increasing their relative population during stress events^43–46^. This is consistent with the ecology of opportunistic pathogens that maintain neutral, commensal or mutualistic interactions with their host but upon sensing host stress, become pathogenic of their host^47,48^. In the Caribbean coral, *Orbicella faveolata,* Vibrionaceae abundance increased in healthy tissue treated with amoxicillin relative to coral without antibiotic treatment^49^. However, this could be due to the loss of amoxicillin sensitive taxa and Vibrionaceae resistance to amoxicillin. Not all *Vibrio* are pathogenic, and the two ASVs could not be resolved to species level^50^. Similarly, marine Gram-positive bacteria, including *Staphylococcus*, are a recognised source of antibacterial secondary metabolites, antibiotic resistance and pathogens^51–53^. Uptake of these ASVs by *Pachyseris* may be associated with compound stress of antibiotic treatment^54^, CMT and environmental perturbations, and not a signature of inoculation success alone, given that they were absent at 10 d.

Two *Endozoicomonas* ASVs had the highest relative abundance and were specific to *Acropora* (Figure 5). They were also both unsuccessful in CMT, being absent in all inoculated treatment groups at 10 d (Figure 5B; Figure 7). *Endozoicomonas* is a ubiquitous marine symbiont, often being dominant ASV(s) in coral microbiomes^55,56^, despite its reported low relative abundance, or absence, in Singaporean corals^57–61^. *Endozoicomonas* is often seen as a marker of coral health and microbiome stability, with specific *Endozoicomonas* strains being promising coral probiotic candidates with strong uptake by a coral host. For example, *E. acroporae* Acr-14^T^, isolated in Kenting, Taiwan, was recently shown to enhance thermal tolerance in *Stylophora pistillata* by mitigating heat-induced protein-folding stress and promoting pro-survival signalling^62^. Some *Endozoicomonas* spp. inhabit within the coral tissue layer, which may reduce their sensitivity to CMT^55,63^. Ruiz-Toquica, et al. ^64^ found *Endozoicomonas* dominance in combination with Vibrionaceae stability to be a marker for *Madracis auretenra* resilience on urban reefs in Colombia. However, we observed low relative abundance of *Endozoicomonas* ASVs in *Pachyseris*, no uptake from CMT, and that its microbiome undergoes high levels of flexibility in their microbiomes overtime (Figure 3), which may reflect an adaptation to the high stress encounter on Singapore’s equatorial turbid reefs^30^.

ASVs successfully transferred via CMT were disproportionately rare and unculturable. ASVs shared between the *Acropora* inoculum and inoculated treatment groups at 10 d (Figure 7) largely corresponded to unculturable environmental lineages, including SAR324 Clade B and SAR11 Clade 1a, rather than dominant donor taxa (e.g. high abundance of *Endozoicomonas* from *Acropora*, Figure 5A). This suggests that CMT can be a successful strategy for unculturable bacteria, whose low abundance reflects a high trophic or keystone position within the community, or low abundance environmental strains may transfer more readily via whole-microbiome inoculation. Such taxa are increasingly recognised as potential keystone members of microbial communities, exerting influence disproportionate to their abundance^65^.

### Towards more targeted probiotic strategies

While coral probiotics can temporarily improve stress tolerance, the best mechanisms of beneficial trait transfer remain unclear^66^. BMC approaches typically focus on one or a small group of selected bacteria^12–14,67^. Thatcher, et al. ^68^ individually inoculated newly settled *Acropora kenti* with pre-screened probiotic strains, where several strains persisted and induced measurable structural change, including microbial aggregates that formed exclusively in *Endozoicomonas*-inoculated corals. The same genus failed to establish in our study when part of a complex and undefined community, with components (e.g. mucus) of the donor, were diluted for the CMT. Thus, our combined results suggest that transplantation success for specific ASVs depends on host specificity, while antibiotic pretreatment would determine overall success in shifting alpha and beta diversity of the recipient microbiome.

A strain specific approach using a *Pseudovibrio* sp. isolated from *Pachyseris speciosa* illustrates how strain-specific probiotics can be: without a ∼490 kbp chromid, this *Pseudovibrio* acts as an opportunistic pathogen that accelerates coral bleaching, but megaplasmid-carrying strains have shifted toward mutualism, producing antibiotics that inhibit *Vibrio* pathogens and conferring a full 1°C increase in coral thermal tolerance^15^. This also shows that probiotic potential here depends on a specific mobile genetic element rather than the species alone.

Other methods, including the use of bacterial carriers, could provide additional tools to increase *in situ* inoculation success^69^. Conversely, some benefits of BMC may not be conferred via direct uptake of the introduced bacteria but rather indirectly conferred via changes to host gene expression and metabolism^13^ or heterotrophic feeding^69,70^. The use of native probiotics is a potentially safe tool to enhance coral stress tolerance *in situ* without negative effects to the surrounding environment^71^. However, the application of BMC and CMT remains a labour-intensive endeavour, often involving multiple rounds of repeated inoculation to achieve a temporary effect, and as shown here, whole-microbiome transplantation does not guarantee reliable or lasting uptake of specific ASVs, even when paired with antibiotic pre-treatment. CMT is compounded by an inherent risk of not being able to guarantee an inoculum free of coral pathogens, and inoculation could plausibly introduce pathogens into the recipient host microbiome alongside beneficial taxa.

Investigations of not just the best probiotics^66^, but also the best methodology to increase the duration of resilience trait inheritance should be prioritized^17,69^, with cultured, strain-specific approaches offering a more tractable and lower-risk path forward than whole-microbiome transplantation^15,62^. Probiotics are proving to be one of the few successful tools for mitigating the negative effects of climate change and environmental stress on coral^72^. Therefore, there is global interest in ensuring it is carried out utilizing the most effective methodology.

## Materials and Methods

### Coral microbiome transplantation experiment

Four *Pachyseris* colonies were collected from the fringing reef off Raffles Lighthouse, Singapore (1°09′39″ N, 103°44′26″ E) and brought to the aquaria facility at St. John’s Island National Marine Laboratory (SJINML). Colonies were fragmented and acclimatized in an outdoor flowthrough tank with sand-filtered seawater (∼ 56 L) for one month before the start of the experiment. Fragments were then rinsed with 0.2 μm filtered seawater (FSW) and moved to indoor 20 L recirculation tanks with UV and ozone treated FSW, pH 8, 31.5-32 ppt salinity. Tanks were submerged in a water bath to maintain a temperature of 29 °C and placed in a room with ambient lighting. There were 12 tanks in total, one tank for each of the four coral colonies across three antibiotic treatments: a control that received no antibiotics, 0.25 mg/mL of the broad spectrum antibiotic ciprofloxacin, and cocktail of ampicillin, nalidixic acid, metronidazole and streptomycin with each 0.05 mg/mL (Figure 1). Following 24 h of antibiotic treatment, half of the fragments were removed from each tank, rinsed with FSW and inoculated with 7 mL suspension of *Acropora* macerate, which were obtained from SJINML coral fragment library. The macerate was freshly prepared using 6 L of high pressured FSW to slough off the tissue of a palm-sized *Acropora* fragment. Inoculated fragments were placed in 5 L FSW for 4 h then individual 1 L outdoor tanks for monitoring, together with uninoculated fragments. Each tank was individually aerated and received flow- through, sand-filtered seawater. Triplicate sampling of each colony and treatment was done at two timepoints: 1 d post microbiome inoculation (1 d) and 10 d post microbiome inoculation (10 d). Coral fragments were snap frozen and transported on dry ice to the Singapore Centre for Environmental Life Sciences Engineering (SCELSE) at Nanyang Technological University (NTU) for DNA extraction, PCR and sequencing of the prokaryotic microbiome community using 16S rDNA amplicon sequencing.

#### DNA extraction, 16S rDNA amplification, sequencing and bioinformatic processing

Following sample collection, coral tissue was separated from all fragments using high pressured sterile water and centrifuged at 12,000 g for 20 min before discarding the supernatant and storing the tissue pellet at -80°C. DNA was extracted using the DNeasy PowerBiofilm Kit with the following modifications adapted from Sunagawa, et al. ^73^: addition of 10 µl lysozyme after adding lysis solution (FB) and a 10 min incubation at room temperature before adding 20 µl of Proteinase K incubation at 65 °C for 1 h. Triplicate blank samples were processed with the same protocol for each of the extraction kits used.

Extracted DNA was stored at -20 °C. PCR reactions in a 20 µl volume were run in triplicate, containing: 10 µl HotStarTaq Plus Master Mix, 1 µl of 10 µM forward and reverse primers,, 5 µl water, 1 µl 100% DMSO, 1 µl 1% BSA, and 1 µl template DNA (5 ng/µl). Primers 515F and 806R were used for amplification of the V4 region of the 16S rRNA gene^74,75^. Thermo- cycling conditions were as follows: initial denaturation at 95 °C for 5 min, followed by 35 cycles of 94 °C for 30 s, 53 °C for 40 s, and 72 °C for 1 min, then a final extension of 10 min at 72 °C. Products were pooled and purified with Agencourt® AMPure XP beads. Purity of the product was checked on an Agilent 2200 TapeStation and quantified with a Qubit™ 2.0 fluorometer. library preparation and amplicon sequencing using 2x300PE on an Illumina MiSeq platform was performed by the Singapore Centre for Environmental Life Sciences Engineering (Nanyang Technological University).

Sequence data were processed following the *Dada2* pipeline version 1.16 to generate amplicon sequence variants (ASVs) for each sample^76^. Briefly, sequence reads were trimmed to 210 bp for the forward read and 190 bp for the reverse read based on visual assessment of sequence read quality, and default filtering parameters were applied (maxN=0, truncQ=2, rm.phix=TRUE and maxEE=2). Error learning algorithms were then applied to the forward and reverse reads, which were merged before removing chimeric sequences. The *decontam* package^77^ utilized the blank extractions to identify and remove contaminating sequence reads. Finally, ASV identity was assigned at genus taxonomic resolution using the built in Bayesian classifier *assignTaxonomy* using SILVA SSU r138.1 as reference library^78^, and any sequences identified as mitochondria, chloroplast or unassigned to a Domain were removed. Samples were rarefied to 13, 222 sequence reads per sample to account for variation in sequencing depths.

### Statistical analyses

Alpha diversity metrics, including observed richness, Chao1, Shannon diversity (H), and inverse Simpson, were calculated using *phyloseq*^79^ and compared by treatment and colony within each timepoint with Kruskal-Wallis tests or ANOVA depending on the significance of Shapiro-Wilk tests for normality. Follow-up Dunn tests for Kruskal-Wallis tests and Tukey’s HSD tests for ANOVA were used for pairwise comparisons if the primary test was significant. Bonferroni corrections were applied to determine significance. Due to differences in sample size, temporal comparisons of alpha diversity between the two time points were tested with Welch’s t-test or the Mann-Whitney U test depending on the significance of Shapiro-Wilk tests for normality.

Beta diversity was compared using permutational multivariate analyses of variance (PERMANOVA). To test whether antibiotic treatments differed from one another within each inoculation status, pairwise PERMANOVA (Bray-Curtis dissimilarity, 999 permutations) was performed between antibiotic treatment groups separately at 1 d and 10 d, split by inoculation status, with p-values adjusted for multiple comparisons using the Benjamini-Hochberg procedure (Table 1, Figure 4).

An indicator species analysis was run with the *multipatt* function in the *indicspecies* package based on a point biserial correlation coefficient function and 9999 permutations^80^. This analysis identifies indicator species based on both ASV relative abundance and fidelity to determine ASVs associated with the selected treatment. This analysis was only applied to samples at 10 d due to insufficient *Acropora* replicates at 1 d. The R package *UpSetR*^29^ was used to determine unique ASVs shared between the *Acropora* microbiome inoculum and the inoculated treatment groups. Potentially inoculated ASVs were compared to *Acropora* microbiome samples collected at the same time point to account for expected changes in the *Acropora* bacterial community over time. Due to quality filtering of the sequence data, there was one replicate sample of inoculum, one replicate of *Acropora* at 1 d, and three replicates of *Acropora* from 10 d. However, *UpSetR* analysis is based on the presence or absence of ASVs, not abundance, so it is less sensitive to uneven replication where beta diversity is low.

## Data availability

Raw sequencing reads were uploaded to the NCBI Sequence Read Archive (SRA) under BioProject PRJNA1479097.

## Acknowledgements

This project is supported by the National Research Foundation, Singapore, and the National Parks Board (NParks), Singapore under its Marine Climate Change Science Programme (MCCS Award NRF-MCCS21-1-5-0001). Any opinions, findings and conclusions or recommendations expressed in this material are those of the author(s) and do not reflect the views of National Research Foundation, Singapore and National Parks Board, Singapore.

The Singapore Centre for Environmental Life Sciences Engineering (SCELSE) is funded by the Ministry of Education, Singapore, the National Research Foundation of Singapore, Nanyang Technological University Singapore (NTU) and National University of Singapore (NUS). The authors would like to acknowledge St. John’s Island National Marine Laboratory (SJINML) for providing the facility necessary for conducting the research. The Laboratory is a National Research Infrastructure under the National Research Foundation Singapore. We would also like to thank the SJI Coral team at the NUS Tropical Marine Science Institute (TMSI) for collecting the coral fragments used in this research (NParks permit NP/RP16- 156-2c).

## Author contributions

LKD and KHP conceived the project and experimental design. LKD and KHP performed the experiments and generated the data. LKD, CWHS and RJC analysed the data. LKD and CWHS contributed to figure preparation. LKD, KHP, CWHS and RJC contributed to writing and editing the manuscript.

## References

1 Rohwer, F., Seguritan, V., Azam, F. & Knowlton, N. Diversity and distribution of coral- associated bacteria. Mar Ecol Prog Ser 243, 1–10 (2002). 10.3354/meps243001

2 Rosenberg, E., Koren, O., Reshef, L., Efrony, R. & Zilber-Rosenberg, I. The role of microorganisms in coral health, disease and evolution. Nat Rev Microbiol 5, 355–362 (2007). 10.1038/nrmicro1635

3 Peixoto, R. S., Rosado, P. M., Leite, D. C. d. A., Rosado, A. S. & Bourne, D. G. Beneficial Microorganisms for Corals (BMC): Proposed mechanisms for coral health and resilience. Front Microbiol 8 (2017). 10.3389/fmicb.2017.00341

4 Bourne, D. G., Morrow, K. M. & Webster, N. S. Insights into the coral microbiome: Underpinning the health and resilience of reef ecosystems. Annu Rev Microbiol 70, 317–340 (2016). 10.1146/annurev-micro-102215-095440

5 Voolstra, C. R. & Ziegler, M. Adapting with microbial help: Microbiome flexibility facilitates rapid responses to environmental change. BioEssays 42, 2000004 (2020). 10.1002/bies.202000004

6 Ziegler, M. et al. Coral bacterial community structure responds to environmental change in a host-specific manner. Nat Commun 10, 3092 (2019). 10.1038/s41467-019-10969-5

7 Ziegler, M., Seneca, F. O., Yum, L. K., Palumbi, S. R. & Voolstra, C. R. Bacterial community dynamics are linked to patterns of coral heat tolerance. Nat Commun 8, 14213 (2017). 10.1038/ncomms14213

8 Zaneveld, J. R. et al. Overfishing and nutrient pollution interact with temperature to disrupt coral reefs down to microbial scales. Nat Commun 7, 11833 (2016). 10.1038/ncomms11833

9. McDevitt-Irwin, J. M., Baum, J. K., Garren, M. & Vega Thurber, R. L. Responses of coral- associated bacterial communities to local and global stressors. Front Mar Sci 4 (2017).

10 Reshef, L., Koren, O., Loya, Y., Zilber-Rosenberg, I. & Rosenberg, E. The coral probiotic hypothesis. Environ Microbiol 8, 2068–2073 (2006). 10.1111/j.1462-2920.2006.01148.x

11 Damjanovic, K., van Oppen, M. J. H., Menéndez, P. & Blackall, L. L. Experimental Inoculation of coral recruits With marine bacteria indicates scope for microbiome manipulation in *Acropora tenuis* and *Platygyra daedalea*. Front Microbiol 10 (2019).

12 Rosado, P. M. et al. Marine probiotics: increasing coral resistance to bleaching through microbiome manipulation. ISME J 13, 921–936 (2019). 10.1038/s41396-018-0323-6

13 Santoro, E. P. et al. Coral microbiome manipulation elicits metabolic and genetic restructuring to mitigate heat stress and evade mortality. Sci Adv 7, eabg3088 (2021). 10.1126/sciadv.abg3088

14 Zhang, Y. et al. Shifting the microbiome of a coral holobiont and improving host physiology by inoculation with a potentially beneficial bacterial consortium. BMC Microbiol 21, 130 (2021). 10.1186/s12866-021-02167-5

15 Sim, C. W. H. et al. Chromid transforms opportunistic pathogen into beneficial symbiont to raise coral’s thermal tipping point. bioRxiv, 2026.2006.2019.733351 (2026). 10.64898/2026.06.19.733351

16 Doering, T. et al. Towards enhancing coral heat tolerance: a “microbiome transplantation” treatment using inoculations of homogenized coral tissues. Microbiome 9, 102 (2021). 10.1186/s40168-021-01053-6

17 Doering, T., Maire, J., van Oppen, M. J. H. & Blackall, L. L. Advancing coral microbiome manipulation to build long-term climate resilience. Microbiol Aust 44, 36–40 (2023). 10.1071/MA23009

18 Sweet, M. J., Croquer, A. & Bythell, J. C. Dynamics of bacterial community development in the reef coral *Acropora muricata* following experimental antibiotic treatment. Coral Reefs 30, 1121–1133 (2011). 10.1007/s00338-011-0800-0

19 Gilbert, J., Hill, R., Doblin, M. & Ralph, P. Microbial consortia increase thermal tolerance of corals. Marine Biology 159, 1763–1771 (2012). 10.1007/s00227-012-1967-9

20 Meron, D. et al. The complexity of the holobiont in the Red Sea coral *Euphyllia paradivisa* under heat stress. Microorganisms 8, 372 (2020).

21 Glasl, B., Herndl, G. J. & Frade, P. R. The microbiome of coral surface mucus has a key role in mediating holobiont health and survival upon disturbance. ISME J 10, 2280–2292 (2016). 10.1038/ismej.2016.9

22 Tun, K. et al. in Status of coral reefs in East Asian Seas region: 2010 9-27 (Global Coral Reef Monitoring Network and Reef and Rainforest Research Center, 2013).

23 Guest, J. R. et al. 27 years of benthic and coral community dynamics on turbid, highly urbanised reefs off Singapore. Sci Rep 6, 36260 (2016). 10.1038/srep36260

24 Ng, C. S. L. & Chou, L. M. Rearing juvenile ‘corals of opportunity’ in in situ nurseries – A reef rehabilitation approach for sediment-impacted environments. Mar Biol Res 10, 833–838 (2014). 10.1080/17451000.2013.853124

25 Guest, J. R. et al. Coral community response to bleaching on a highly disturbed reef. Sci Rep 6, 20717 (2016). 10.1038/srep20717

26 Wong, J. S. Y. et al. Comparing patterns of taxonomic, functional and phylogenetic diversity in reef coral communities. Coral Reefs 37, 737–750 (2018). 10.1007/s00338-018-1698-6

27 Feldman, B. et al. Distinct lineages and population genomic structure of the coral *Pachyseris speciosa* in the small equatorial reef system of Singapore. Coral Reefs 41 (2021). 10.1007/s00338-021-02160-4

28 Chow, G. S. E., Chan, Y. K. S., Jain, S. S. & Huang, D. Light limitation selects for depth generalists in urbanised reef coral communities. Mar Environ Res 147, 101–112 (2019). 10.1016/j.marenvres.2019.04.010

29 Conway, J. R., Lex, A. & Gehlenborg, N. UpSetR: an R package for the visualization of intersecting sets and their properties. Bioinformatics 33, 2938–2940 (2017). 10.1093/bioinformatics/btx364

30 Guest, J. R. et al. Contrasting patterns of coral bleaching susceptibility in 2010 suggest an adaptive response to thermal stress. PLoS ONE 7, e33353 (2012). 10.1371/journal.pone.0033353

31 Penin, L., Vidal-Dupiol, J. & Adjeroud, M. Response of coral assemblages to thermal stress: are bleaching intensity and spatial patterns consistent between events? Environ Monit Assess 185, 5031–5042 (2013). 10.1007/s10661-012-2923-3

32 Evensen, N. R., Fine, M., Perna, G., Voolstra, C. R. & Barshis, D. J. Remarkably high and consistent tolerance of a Red Sea coral to acute and chronic thermal stress exposures. Limnol Oceanog 66, 1718–1729 (2021). 10.1002/lno.11715

33 Marshall, P. A. & Baird, A. H. Bleaching of corals on the Great Barrier Reef: differential susceptibilities among taxa. Coral Reefs 19, 155–163 (2000). 10.1007/s003380000086

34 Loya, Y. et al. Coral bleaching: the winners and the losers. Ecology Letters 4, 122–131 (2001). 10.1046/j.1461-0248.2001.00203.x

35 van Oppen, M. J. H. et al. Shifting paradigms in restoration of the world’s coral reefs. Global Change Biology 23, 3437–3448 (2017). 10.1111/gcb.13647

36 Howells, E. J. et al. Enhancing the heat tolerance of reef-building corals to future warming. Sci Adv 7, eabg6070 (2021). 10.1126/sciadv.abg6070

37 Humanes, A. et al. Selective breeding enhances coral heat tolerance to marine heatwaves. Nat Commun 15, 8703 (2024). 10.1038/s41467-024-52895-1

38 Quigley, K. M. Breeding and selecting corals resilient to global warming. Annu Rev Anim Biosci 12, 209–332 (2024). 10.1146/annurev-animal-021122-093315

39 Pollock, F. J. et al. Coral-associated bacteria demonstrate phylosymbiosis and cophylogeny. Nat Commun 9, 4921 (2018). 10.1038/s41467-018-07275-x

40 Kohanski, M. A., Dwyer, D. J. & Collins, J. J. How antibiotics kill bacteria: from targets to networks. Nat Rev Microbiol 8, 423–435 (2010). 10.1038/nrmicro2333

41 Newton, D. P., Ho, P.-Y. & Huang, K. C. Modulation of antibiotic effects on microbial communities by resource competition. Nat Commun 14, 2398 (2023). 10.1038/s41467-023-37895-x

42 Roemhild, R., Bollenbach, T. & Andersson, D. I. The physiology and genetics of bacterial responses to antibiotic combinations. Nat Rev Microbiol 20, 478–490 (2022). 10.1038/s41579-022-00700-5

43 Munn, C. B. The role of *Vibrios* in diseases of corals. Microbiol Spectr 3, 10.1128/microbiolspec.ve-0006-2014 (2015). 10.1128/microbiolspec.ve-0006-2014

44 Tout, J. et al. Increased seawater temperature increases the abundance and alters the structure of natural *Vibrio* populations associated with the coral *Pocillopora damicornis*. Front Microbiol 6 (2015).

45 Divya, S., Thinesh, T., Seghal Kiran, G., Hassan, S. & Selvin, J. Emergence of a multi host biofilm forming opportunistic pathogen *Staphylococcus sciuri* D26 in coral *Favites abdita*. Microb Pathog 120, 204–212 (2018). 10.1016/j.micpath.2018.04.037

46 Luk, H. C. et al. Draft genomes of two *Vibrio* spp. isolated from the brown alga *Sargassum ilicifolium* in Singapore. Microbiol Resour Announc 14, e00328–00325 (2025). 10.1128/mra.00328-25

47 Kushmaro, A., Loya, Y., Fine, M. & Rosenberg, E. Bacterial infection and coral bleaching [6]. Nature 380, 396–396 (1996). 10.1038/380396a0

48 Bramucci, A. et al. The bacterial symbiont *Phaeobacter inhibens* shapes the life history of its algal host *Emiliania huxleyi*. Front Mar Sci 5, 188 (2018). 10.3389/fmars.2018.00188

49 Pearson-Lund Alexi, S., et al. Evaluating the effect of amoxicillin treatment on the microbiome of *Orbicella faveolata* with Caribbean yellow band disease. Appl Environ Microbiol 91, e02407–02424 (2025). 10.1128/aem.02407-24

50 Cano-Gomez, A., Høj, L., Owens, L. & Andreakis, N. Multilocus sequence analysis provides basis for fast and reliable identification of *Vibrio harveyi*-related species and reveals previous misidentification of important marine pathogens. Syst Appl Microbiol 34, 561–565 (2011). 10.1016/j.syapm.2011.09.001

51 Gontang, E. A., Fenical, W. & Jensen, P. R. Phylogenetic diversity of gram-positive bacteria cultured from marine sediments. Appl Environ Microbiol 73, 3272–3282 (2007). 10.1128/AEM.02811-06

52 Fenical, W. & Jensen, P. R. Developing a new resource for drug discovery: marine actinomycete bacteria. Nat Chem Biol 2, 666–673 (2006). 10.1038/nchembio841

53 Maithani, P. et al. Staphylococcaceae isolates from the reef macroalga *Sargassum ilicifolium*, found in the coastal waters of Singapore. Microbiol Resour Announc 14, e00335–00325 (2025). 10.1128/mra.00335-25

54 Patton, S. et al. Antibiotic type and dose variably affect microbiomes of a disease-resistant *Acropora cervicornis* genotype. Environmental Microbiome 20, 46 (2025). 10.1186/s40793-025-00709-2

55 Neave, M. J. et al. Differential specificity between closely related corals and abundant *Endozoicomonas* endosymbionts across global scales. ISME J 11, 186–200 (2017). 10.1038/ismej.2016.95

56 Pogoreutz, C. et al. Coral holobiont cues prime *Endozoicomonas* for a symbiotic lifestyle. ISME J 16, 1883–1895 (2022). 10.1038/s41396-022-01226-7

57 Wainwright, B. J., Afiq-Rosli, L., Zahn, G. L. & Huang, D. Characterisation of coral- associated bacterial communities in an urbanised marine environment shows strong divergence over small geographic scales. Coral Reefs 38, 1097–1106 (2019). 10.1007/s00338-019-01837-1

58 Fong, J. et al. Contact- and water-mediated effects of macroalgae on the physiology and microbiome of three Indo-pacific coral species. Front Mar Sci 6 (2020).

59 Deignan, L. K. & McDougald, D. Differential response of the microbiome of *Pocillopora acuta* to reciprocal transplantation within Singapore. Microbial Ecology 83, 608–618 (2022). 10.1007/s00248-021-01793-w

60 Moynihan, M. A. et al. Coral-associated nitrogen fixation rates and diazotrophic diversity on a nutrient-replete equatorial reef. ISME J 16, 233–246 (2022). 10.1038/s41396-021-01054-1

61 Deignan, L. K., Pwa, K. H., Loh, A. A. R., Rice, S. A. & McDougald, D. The microbiomes of two Singaporean corals show site-specific differentiation and variability that correlates with the seasonal monsoons. Coral Reefs 42, 677–691 (2023). 10.1007/s00338-023-02376-6

62 Lu, C.-Y., et al. *Endozoicomonas acroporae* enhances coral thermal resilience through host– microbe coordination. *ISME J*, wrag173 (2026). 10.1093/ismejo/wrag173

63 Bayer, T. et al. The microbiome of the Red Sea coral *Stylophora pistillata* is dominated by tissue-associated *Endozoicomonas* bacteria. Appl Environ Microbiol 79, 4759–4762 (2013). 10.1128/AEM.00695-13

64 Ruiz-Toquica, J., Franco Herrera, A. & Medina, M. *Endozoicomonas* dominance and Vibrionaceae stability underpin resilience in urban coral *Madracis auretenra*. PeerJ 13, e19226 (2025). 10.7717/peerj.19226

65 Banerjee, S., Schlaeppi, K. & van der Heijden, M. G. A. Keystone taxa as drivers of microbiome structure and functioning. Nat Rev Microbiol 16, 567–576 (2018). 10.1038/s41579-018-0024-1

66 Rosado, P. M. et al. Exploring the potential molecular mechanisms of interactions between a probiotic consortium and its coral host. mSystems 8, e00921–00922 (2023). 10.1128/msystems.00921-22

67 Fragoso ados Santos, H., et al. Impact of oil spills on coral reefs can be reduced by bioremediation using probiotic microbiota. Sci Rep 5, 18268 (2015). 10.1038/srep18268

68 Thatcher, C. et al. Bacterial dynamics in newly settled *Acropora kenti*: Insights from inoculations with individual probiotic candidates. Environ Microbiol 27, e70143 (2025). 10.1111/1462-2920.70143

69 Assis, J. M. et al. Delivering beneficial microorganisms for corals: Rotifers as carriers of probiotic bacteria. Front Microbiol 11 (2020).

70 Martinez, S., Grover, R. & Ferrier-Pagès, C. Unveiling the importance of heterotrophy for coral symbiosis under heat stress. mBio 15, e01966–01924 (2024). 10.1128/mbio.01966-24

71 Delgadillo-Ordoñez, N. et al. Probiotics reshape the coral microbiome in situ without detectable off-target effects in the surrounding environment. Commun Biol 7, 434 (2024). 10.1038/s42003-024-06135-3

72 Peixoto, R. S. et al. Harnessing the microbiome to prevent global biodiversity loss. Nat Microbiol 7, 1726–1735 (2022). 10.1038/s41564-022-01173-1

73 Sunagawa, S., Woodley, C. M. & Medina, M. Threatened corals provide underexplored microbial habitats. PLoS ONE 5, e9554 (2010). 10.1371/journal.pone.0009554

74 Apprill, A., McNally, S., Parsons, R. & Weber, L. Minor revision to V4 region SSU rRNA 806R gene primer greatly increases detection of SAR11 bacterioplankton. Aquat Microb Ecol 75, 129–137 (2015). 10.3354/ame01753

75 Parada, A. E., Needham, D. M. & Fuhrman, J. A. Every base matters: assessing small subunit rRNA primers for marine microbiomes with mock communities, time series and global field samples. Environ Microbiol 18, 1403–1414 (2016). 10.1111/1462-2920.13023

76 Callahan, B. J. et al. DADA2: High-resolution sample inference from Illumina amplicon data. Nat Methods 13, 581–583 (2016). 10.1038/nmeth.3869

77 Davis, N. M., Proctor, D. M., Holmes, S. P., Relman, D. A. & Callahan, B. J. Simple statistical identification and removal of contaminant sequences in marker-gene and metagenomics data. Microbiome 6, 226 (2018). 10.1186/s40168-018-0605-2

78 Quast, C. et al. The SILVA ribosomal RNA gene database project: improved data processing and web-based tools. Nucleic Acids Res 41, D590–596 (2013). 10.1093/nar/gks1219

79 McMurdie, P. J. & Holmes, S. *Phyloseq*: An R package for reproducible interactive analysis and graphics of microbiome census data. PLoS ONE 8, e61217 (2013). 10.1371/journal.pone.0061217

80 Cáceres, M. D. & Legendre, P. Associations between species and groups of sites: indices and statistical inference. Ecology 90, 3566–3574 (2009). 10.1890/08-1823.1

