## Supplementary Material for "Successful coral microbiome transplant from high to low heat tolerant corals requires antibiotic pretreatment"

Running title: Coral microbiome transplantation

**ORCid**

Lindsey Kane DEIGNAN: 0000-0002-5189-0707

Clarence Wei Hung SIM: 0000-0003-2190-7261

Keay Hoon PWA: NA

Rebecca Josephine CASE: 0000-0003-2417-2969

### Supplementary Tables

**Table S1.** Mean alpha diversity metrics (± standard deviation) by treatment group and *Pachyseris* colony Day 1 and Day 10 post inoculation with *Acropora* microbiome. Bold values represent significant (<0.05) temporal comparisons between Day 1 and Day 10. An asterisk (*) is used to indicate significant (<0.05) within timepoint comparisons for treatment and colony. CIP = ciprofloxacin; Inoc = Acropora inoculation; Ctrl = control; mix = antibiotic cocktail.

| | **Day 1 Post Inoculation** | | | | **Day 10 Post Inoculation** | | | |
|---|---|---|---|---|---|---|---|---|
| - | Observed | Chao1 | Shannon | InvSimpson | Observed | Chao1 | Shannon | InvSimpson |
| *Treatment* | | | | | | | | |
| CIP | 442.75 ± 143.44 | 447.03 ± 145.76 | 5.43 ± 0.39 | 130.90 ± 53.68 | 525.83 ± 170.52 | 531.98 ± 176.77 | 5.46 ± 0.39 | 120.89 ± 65.88 |
| CIP_Inoc | 482.00 ± 283.96 | 490.27 ± 293.91 | **5.16 ± 0.60** | **77.85 ± 52.67** | 671.67 ± 255.08 | 690.67 ± 277.46 | **5.70 ± 0.38** | **151.38 ± 62.50** |
| Ctrl | 485.86 ± 291.91 | 495.37 ± 300.64 | 5.11 ± 0.75 | 86.89 ± 49.96 | 520.08 ± 216.78 | 530.82 ± 227.88 | 5.12 ± 0.69 | 79.44 ± 52.95 |
| Ctrl_Inoc | 567.50 ± 239.53 | 584.52 ± 247.33 | 5.00 ± 0.58 | 59.88 ± 47.04 | 607.83 ± 228.73 | 618.10 ± 233.62 | 5.35 ± 0.82 | 101.42 ± 61.95 |
| MIX | 433.25 ± 277.13 | 438.04 ± 281.65 | 4.91 ± 1.42 | 114.65 ± 101.24 | 540.50 ± 238.45 | 546.18 ± 242.11 | 5.36 ± 0.68 | 121.05 ± 81.80 |
| MIX_Inoc | 390.88 ± 221.23 | 396.81 ± 227.08 | **4.95 ± 0.52** | **63.36 ± 26.39** | 544.75 ± 194.12 | 553.50 ± 201.91 | **5.50 ± 0.38** | **121.53 ± 54.18** |
| *Colony* | | | | | | | | |
| *Pachyseris*_1 | 713.88 ± 314.90* | 729.26 ± 324.70* | 5.60 ± 0.67 | 115.85 ± 61.42 | 656.67 ± 214.58 | 671.87 ± 230.02 | 5.57 ± 0.59 | 128.93 ± 64.91 |
| *Pachyseris*_2 | 505.18 ± 197.58 | 514.40 ± 202.79 | 5.25 ± 0.52 | 103.28 ± 82.75 | 615.67 ± 257.14 | 628.96 ± 266.68 | 5.44 ± 0.69 | 112.04 ± 59.68 |
| *Pachyseris*_3 | **339.00 ± 196.92*** | **342.14 ± 200.77*** | **4.69 ± 1.01** | **64.81 ± 41.73** | **486.33 ± 179.28** | **491.82 ± 183.73** | **5.31 ± 0.53** | **116.21 ± 67.82** |
| *Pachyseris*_4 | **388.22 ± 127.76** | 396.85 ± 136.73 | 4.96 ± 0.40 | 71.75 ± 51.10 | **515.11 ± 183.80** | 521.53 ± 190.90 | 5.33 ± 0.56 | 106.63 ± 72.35 |
| *Acropora* | n/a | n/a | n/a | n/a | 652.67 ± 264.95 | 682.39 ± 270.93 | 5.11 ± 0.58 | 55.25 ± 19.51 |
| Total | **464.48 ± 247.32** | **472.72 ± 254.66** | **5.07 ± 0.79** | **86.09 ± 61.87** | **571.81 ± 218.76** | **582.70 ± 228.87** | **5.40 ± 0.59** | **113.52 ± 65.31** |

**Table S2.** List of ASVs identified as indicators of *Acropora* and their mean abundance by treatment group. An asterisk (*) with bolded taxa indicates the four ASV whose mean abundance is higher in the inoculation group compared to the non-inoculation group.

| ASV | Phylum | Lowest taxonomic rank | Acropora | Ctrl | Ctrl_Inoc | CIP | CIP_Inoc | MIX | MIX_Inoc |
|---|---|---|---|---|---|---|---|---|---|
| ASV48 | Proteobacteria | Genus *Endozoicomonas* | 417.33 | 0.00 | 0.00 | 0.00 | 0.00 | 0.00 | 0.00 |
| ASV73 | Proteobacteria | Genus *Endozoicomonas* | 289.33 | 0.00 | 0.00 | 0.00 | 0.00 | 0.00 | 0.00 |
| ASV111 | Proteobacteria | Genus *Alteromonas* | 39.67 | 0.67 | 3.08 | 13.00 | 3.08 | 7.42 | 0.75 |
| ASV120 | Proteobacteria | Genus *Pyruvatibacter* | 81.33 | 7.00 | 12.50 | 59.92 | 23.67 | 2.50 | 28.67 |
| ASV134 | Proteobacteria | Genus Marine Methylotrophic G3 | 556.33 | 2.25 | 2.08 | 7.25 | 1.58 | 20.33 | 4.25 |
| ASV162 | Proteobacteria | Family Alteromonadaceae | 583.67 | 2.92 | 3.58 | 12.17 | 4.25 | 0.00 | 2.17 |
| ASV168 | Proteobacteria | Genus Marine Methylotrophic G3 | 447.33 | 2.58 | 1.92 | 4.67 | 1.42 | 14.92 | 4.83 |
| ASV199 | Proteobacteria | Family Rhodobacteraceae | 27.67 | 0.92 | 2.83 | 4.58 | 1.00 | 18.00 | 9.50 |
| ASV212 | Proteobacteria | Family Alteromonadaceae | 446.33 | 1.67 | 3.33 | 8.17 | 2.50 | 0.00 | 2.42 |
| ASV259 | Proteobacteria | Family Rhodobacteraceae | 19.33 | 1.25 | 0.42 | 7.75 | 0.33 | 12.17 | 7.25 |
| ASV261 | Proteobacteria | Genus *Ruegeria* | 49.33 | 5.67 | 2.75 | 37.58 | 4.92 | 22.00 | 24.25 |
| ASV335 | Proteobacteria | Genus *Thalassolituus* | 246.00 | 29.17 | 2.42 | 0.00 | 1.08 | 0.00 | 0.92 |
| **ASV349*** | **Proteobacteria** | **Genus *Shimia*** | 78.67 | 2.00 | 7.42 | 11.58 | 8.25 | 9.58 | 22.67 |
| ASV448 | Proteobacteria | Genus *Thalassolituus* | 209.33 | 22.58 | 2.67 | 1.00 | 1.08 | 0.00 | 0.42 |
| ASV458 | Planctomycetota | Genus *Blastopirellula* | 46.33 | 0.42 | 7.75 | 6.33 | 10.25 | 17.67 | 5.33 |
| **ASV494*** | **Proteobacteria** | **Genus *Shimia*** | 58.67 | 0.00 | 6.58 | 11.00 | 1.92 | 4.42 | 14.25 |
| ASV734 | Proteobacteria | Genus HTCC5015 | 70.00 | 1.33 | 0.00 | 7.42 | 0.00 | 2.92 | 7.83 |
| **ASV871*** | **Proteobacteria** | **Genus *Marinicella*** | 16.33 | 0.58 | 2.58 | 8.00 | 5.58 | 2.58 | 6.83 |
| ASV880 | Proteobacteria | Genus *Oleiphilus* | 19.00 | 1.25 | 0.83 | 0.83 | 0.92 | 3.75 | 2.17 |
| ASV1108 | Bacteroidota | Genus *Aquibacter* | 64.67 | 0.42 | 0.00 | 0.58 | 0.00 | 0.00 | 0.00 |
| ASV1300 | Planctomycetota | Class OM190 | 14.67 | 2.50 | 1.25 | 2.17 | 1.17 | 0.67 | 1.83 |
| ASV1747 | Proteobacteria | Family Methylophagaceae | 45.67 | 0.00 | 0.00 | 2.08 | 0.00 | 0.00 | 0.00 |
| ASV2177 | Planctomycetota | Class BD7-11 | 24.00 | 0.42 | 0.00 | 0.42 | 0.17 | 0.00 | 0.00 |
| ASV2260 | Planctomycetota | Class BD7-11 | 27.33 | 0.00 | 0.00 | 2.00 | 0.00 | 0.00 | 1.25 |
| ASV2515 | Proteobacteria | Family Methylophagaceae | 34.67 | 0.00 | 0.00 | 0.75 | 0.00 | 0.00 | 0.00 |
| ASV2627 | Proteobacteria | Order Rhodospirillales | 36.00 | 0.00 | 0.00 | 1.75 | 0.00 | 0.00 | 0.00 |
| **ASV2637*** | **Planctomycetota** | **Genus *Fuerstia*** | 14.67 | 0.00 | 3.67 | 1.08 | 0.17 | 0.00 | 1.92 |
| ASV4231 | Proteobacteria | Family PS1 clade | 10.33 | 0.42 | 0.00 | 2.67 | 1.83 | 0.58 | 0.00 |
| ASV4261 | Verrucomicrobiota | Family PRD18C08 | 9.33 | 0.00 | 0.08 | 3.50 | 0.00 | 2.75 | 0.58 |
| ASV4563 | Bacteroidota | Genus *Vicingus* | 16.00 | 0.42 | 0.00 | 1.75 | 0.00 | 0.00 | 0.00 |

**Table S3**. The ASVs at Day 1 and Day 10 that were uniquely present in the inoculum and inoculated treatment groups (ASVs reported as mean abundance by treatment group).

| | | Phylum | Lowest taxonomic rank | Ctrl_Inoc | CIP_Inoc | MIX_Inoc | Inoculum | Day 1 *Acropora* | Day 10 *Acropora* |
|---|---|---|---|---|---|---|---|---|---|
| *Day 1* | ASV546 | Proteobacteria | Genus *Vibrio* | 5.5 | 4.89 | 7.63 | 8 | 58 | |
| | ASV908 | Proteobacteria | Genus *Vibrio* | 2.5 | 3 | 5.25 | 5 | 38 | |
| | ASV1597 | Firmicutes | Genus *Staphylococcus* | 0 | 0.33 | 2.25 | 28 | 0 | |
| | ASV6765 | Proteobacteria | Order Defluviicoccales | 0 | 2.67 | 0 | 3 | 0 | |
| *Day 10* | ASV387 | SAR324 clade(Marine group B) | Phylum SAR324 clade (Marine group B) | 0.17 | 0 | 0 | 5 | | 0 |
| | ASV439 | Proteobacteria | Genus Clade Ia | 0 | 4.5 | 0 | 89 | | 0 |
| | ASV590 | Proteobacteria | Genus Clade Ia | 0 | 3.58 | 0 | 43 | | 0 |
| | ASV596 | Proteobacteria | Genus *Neptuniibacter* | 0 | 1.42 | 0 | 3 | | 5 |
| | ASV897 | Proteobacteria | Genus *Massilia* | 1.75 | 0 | 0 | 242 | | 0 |
| | ASV1253 | Proteobacteria | Genus *Massilia* | 1.33 | 0 | 0 | 156 | | 0 |
| | ASV1793 | Bacteroidota | Genus *Cloacibacterium* | 1.58 | 0 | 0 | 9 | | 0 |
| | ASV2008 | Firmicutes | Genus *Staphylococcus* | 0.83 | 0 | 0 | 20 | | 0 |
